# Cell-type-specific alternative polyadenylation (APA) genes reveal the function of dynamic APA in complex tissues

**DOI:** 10.1101/2020.07.30.229096

**Authors:** Yulong Bai, Yidi Qin, Zhenjiang Fan, Robert M. Morrison, KyongNyon Nam, Hassane Mohamed Zarour, Radosveta Koldamova, Quasar Saleem Padiath, Soyeon Kim, Hyun Jung Park

**Affiliations:** Department of Human Genetics, Graduate School of Public Health, University of Pittsburgh, Pittsburgh, USA; Department of Computer Science, School of Computing and Information, University of Pittsburgh, Pittsburgh, USA; Department of Environmental and Occupational Health, Graduate school of Public Health, University of Pittsburgh, Pittsburgh, USA; Department of Neurobiology, School of Medicine, University of Pittsburgh, Pittsburgh, USA; Department of Medicine and Division of Hematology/Oncology, University of Pittsburgh, School of Medicine, Pittsburgh, USA; Department of Immunology, University of Pittsburgh, School of Medicine, Pittsburgh, USA; Department of Computational and Systems Biology, University of Pittsburgh Medical Center, Pittsburgh, USA; Department of Pediatrics, University of Pittsburgh Medical Center, Pittsburgh, USA; Division of Pulmonary Medicine, Children’s Hospital of Pittsburgh of UPMC, Pittsburgh, Pennsylvania, USA

**Keywords:** post-transcriptional regulation, alternative polyadenylation, single-cell RNA

## Abstract

Alternative polyadenylation (APA) causes shortening or lengthening of the 3’-untranslated region (3’-UTR) of genes across multiple cell types. Bioinformatic tools have been developed to identify genes that are affected by APA (APA genes) in single-cell RNA-Seq (scRNA-Seq) data. However, they suffer from low power, and they cannot identify APA genes specific to each cell type (cell-type-specific APA) when multiple cell types are analyzed. To address these limitations, we developed scMAPA that systematically integrates two novel steps. First, scMAPA quantifies 3’-UTR long and short isoforms without requiring assumptions on the read density shape of input data. Second, scMAPA estimates the significance of the APA genes for each cell type while controlling confounders. In the analyses on our novel simulation data and human peripheral blood mono cellular data, scMAPA showed enhanced power in identifying APA genes. Further, in mouse brain data, scMAPA identifies cell-type-specific APA genes, improving interpretability for the cell-type-specific function of APA. We further showed that this improved interpretability helps to understand a novel role of APA on the interaction between neurons and blood vessels, which is critical to maintaining the operational condition of brains. With high sensitivity and interpretability, scMAPA shed novel insights into the function of dynamic APA in complex tissues.

**Key Points:**

- We developed a bioinformatic tool, scMAPA, that identifies dynamic APA across multiple cell types and a novel simulation pipeline to assess performance of such tools in APA calling.
- In simulation data of various scenarios from our novel simulation pipeline, scMAPA achieves sensitivity with a minimal loss of specificity.
- In human peripheral blood monocellular data, scMAPA identifies APA genes accurately and robustly, finding unique associations of APA with hematological processes.
- scMAPA identifies APA genes specific to each cell type in mouse brain data while controlling confounders that sheds novel insights into the complex molecular processes.

## INTRODUCTION

The majority of mammalian messenger RNAs contain multiple polyadenylation (pA) sites, such as proximal and distal, in their 3’-untranslated region (3’-UTR) ^1,2^. By transcribing with various pA sites, alternative polyadenylation (APA) produces distinct isoforms with various lengths of the 3’-UTRs (long and short 3’-UTR isoforms using distal and proximal pA sites, respectively). These APA events are involved in diverse physiological and pathological processes ^3^. For example, global 3’-UTR shortening events promote tumorigenesis by removing microRNA binding sites in several types of cancer^4–6^. Notably, these events occur in tissue-specific and cell-type-specific manners ^1–7^. To identify genes with APA events (APA genes) specific to each cell type (cell-type-specific APA genes), analyzing single-cell RNA sequencing (scRNA-Seq) data is essential, since the data present transcriptome of various cell types.

For scRNA-Seq data, several tools have been developed to identify APA genes across cell types (dynamic APA genes), such as scDAPA^8^, Sierra ^9^ and scAPA^10^. While they have different strengths in identifying dynamic APA, several limitations remain to identify cell-type-specific APA genes in scRNA-Seq data. First, they are based on assumptions on the input read density shape: since several scRNA-Seq utilize 3’ selection and enrichment steps in library construction, accumulation of the reads that originate from a common pA site forms a peak. Based on that, the APA identification methods assumed that signal shapes are different from noise in their peak calling. However, these assumptions are not guaranteed to hold for all genes. For example, for FLT3, a critical gene whose abnormality leads to blood disorders ^11^, 3’ tags form peaks with different shapes and lengths between pDC/HSPC and B/NK cells (**S. Fig. 1A**) in the scRNA-Seq data on Peripheral Blood Monocellular Cells (PBMC) of a healthy donor (10k in https://www.10xgenomics.com/), complicating the quantification process. Due to the reason, with existing methods, FLT3 can hardly be detected.

Second, APA genes cannot be identified for each cell cluster in multi-cluster settings, even though it is critical to study the impact of APA events for each cell type. scDAPA and Sierra identify APA genes mainly between two cell clusters and are not directly applicable for more than two clusters. While scAPA is the only method to identify APA genes in more than two clusters, it statistically tests if the APA usage (the ratio of long and short 3’-UTR isoforms) of each gene is similar across cell clusters and does not estimate the statistical significance in each cell cluster.

Third, the existing methods do not control confounding factors, which is critical to identify APA genes from complex tissues. Confounding arises when cells are affected by factors that are not parts of the research hypothesis under investigation. For example, since brain transcriptome is known to be specific to regions (e.g. cortex and dorsal midbrain) and cell types (e.g. neuron and astrocyte) ^12–14^, one may need to control brain region as confounders depending on how brain cells are clustered and the research question.

Fourth, there is no simulation platform to compare statistical power and specificity of the methods that identify dynamic APA in scRNA-Seq data. Although such a platform is necessary to evaluate those methods with ground truths, it has been challenging since it is not clear how the read density shapes differ between APA and non-APA genes.

To address the first limitation and identify dynamic APA without the assumptions on the read density shape, we hypothesize that formulating an optimization problem instead of a signal processing problem could greatly increase the power since the optimization problem can consider those genes that do not fit the assumptions required by signal processing steps. For the second and the third limitation, we hypothesize that a modeling of each transcript with covariates explicitly representing the cell clusters and all confounders could determine cell-type-specific APA genes while controlling confounders. For the fourth limitation, we hypothesize that the read density shape mainly consists of the ratio of 3’-UTR long and short isoforms.

scMAPA systematically incorporates solving the optimization problem and building the statistical model. Using our novel simulation data and human peripheral blood mono cellular data, we will show how scMAPA enhances statistical power in identifying APA genes. Further, using mouse brain data, we will demonstrate how scMAPA facilitates to understand cell-type-specific functional relevance of APA in a statistically rigorous way.

## RESULTS

### Alternative Polyadenylation identification across multiple cell groups of single-cell RNA-Seq data (scMAPA)

To identify APA genes in scRNA-Seq data without assumptions on the read density shape, scMAPA firstly quantifies 3’UTR long and short isoforms without such assumptions. Using the fact that each 3’ biased read represents the most 3’ end part of a transcript, scMAPA pads each read along the 3’UTR region from the 3’UTR start site to where the read ends (step 1 in **Fig. 1**, see Methods). This transformation enables to compare the ratio of 3’UTR long and short isoforms among cell types without assumptions on the read density shape. For example, this transformation reveals different 3’UTR isoform ratios of FLT3 between pDC/HSPC and B/NK cells in the PBMC data (**S. Fig. 1B**). Due to this transformation, inferring the proximal pA site, a critical step to quantify 3’UTR isoforms, becomes an optimization problem of minimizing the difference between the accumulated density shape of the quantified isoforms and the input read density (step 2 in **Fig. 1)**. Since the difference can be calculated by a quadratic function, we incorporated quadratic programming^15^ to identify inferring the proximal pA site. To apply quadratic programming to identify APA genes in scRNA-Seq data, we extended multiple modules of DaPars ^16^. DaPars incorporated quadratic programming approach to identify APA genes in bulk RNA-Seq data, although it is not directly applicable for multi-cluster settings.

**Figure 1.**
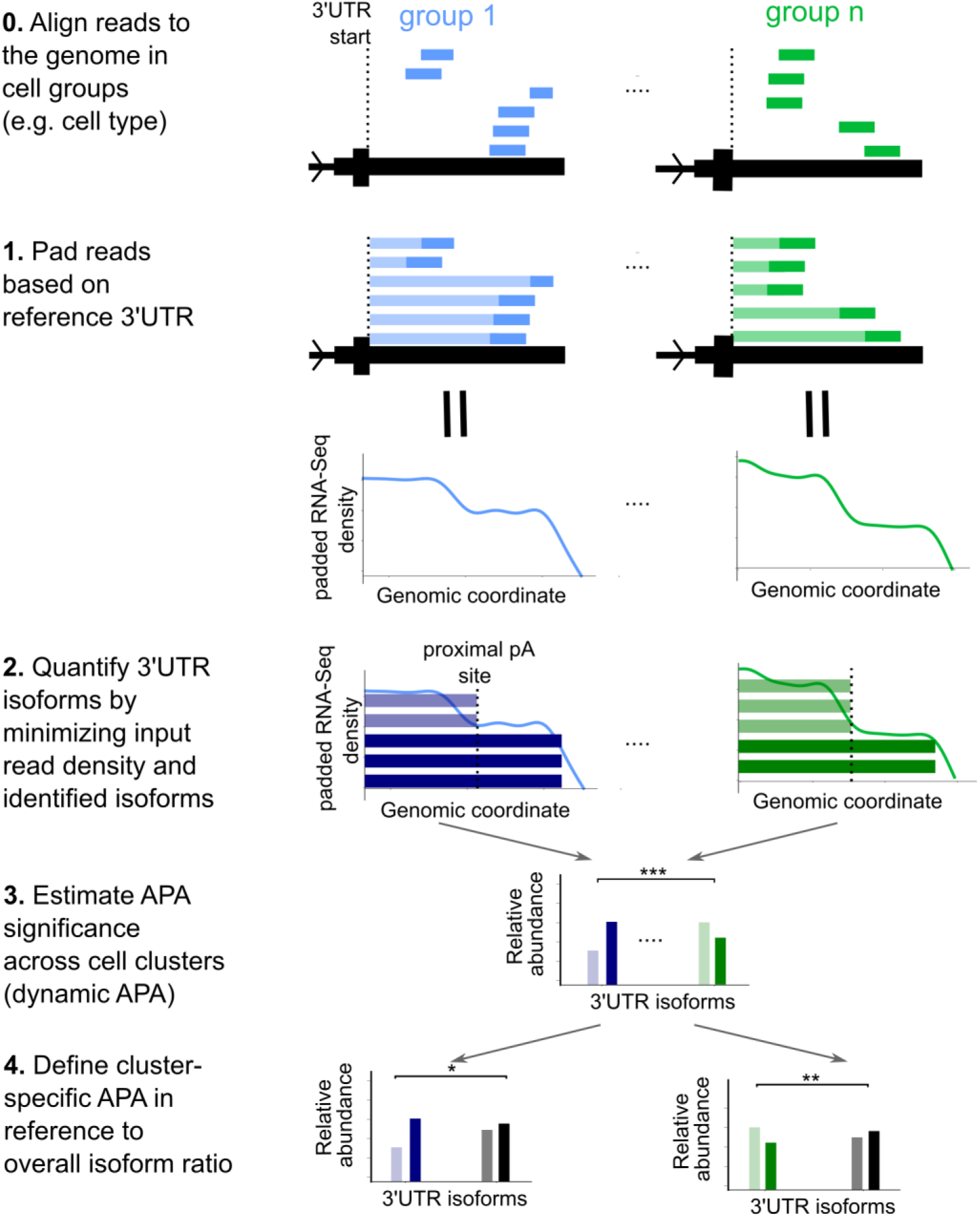
Schematic illustration of each step of scMAPA. In Step 0 and 1, bars in solid color represent 3’ biased scRNA-Seq reads and bars in light color indicate how the 3’ biased reads are padded from the 3’ start site to the end of the read to represent the full-length 3’ UTR of the transcript. In Step 2, bars in dark blue and green indicate the estimated isoforms in each cell type, where solid and light coloring mode indicate 3’ UTR long and short isoforms. In Step 3 and 4, the bars represent the total number of isoforms in each case. The black bars on the bottom represent the grand mean of all long/short isoforms across the groups.

To identify APA genes in each cell type based on the quantification above, scMAPA develops an additive regression model. While the quantified isoforms can be directly used to identify dynamic APA (APA genes across cell types, step 3 in **Fig. 1**), scMAPA further identifies cell-type-specific APA genes (APA genes specific to each cell type, step 4 in **Fig. 1**) by developing a model using each of the isoforms with cell type indicators (see Methods). Since this modeling is flexible to additionally incorporate potential confounders, such as tissue type, age, or sex, scMAPA can control such confounders.

### scMAPA identifies true APA events with an enhanced statistical power

To compare statistical power of scMAPA and scAPA with ground truth APA events, we simulated 3’-UTR long and short isoform expressions for APA and non-APA genes. First, we learned simulation parameters in the mouse brain scRNA-Seq data where five main cell types are defined (neurons, astrocytes, immune cells, oligodendrocytes and vascular) ^17^. In the data, we determined APA and non-APA genes as those detected by both scAPA, evaluated the proportion of the long and short isoforms in the APA and non-APA genes respectively, and calculated standard deviation (SD) of the proportions (*SD_isoprop_*) across the five cell types (see Methods, **S. Fig. 2A)**. The APA genes have wider distributions of the 3’-UTR isoform proportions across the cell types than non-APA genes (0.127 vs. 0.009 on average in terms of *SD_isoprop_*, **S. Fig. 2B**). Second, using *SD_isoprop_* for APA and non-APA genes as simulation parameters, we generated the 3’-UTR isoform proportions for each of 500 APA and 4,500 non-APA genes across 5 groups, each of 600 cells. These isoform proportion values simulated for APA and non-APA genes were further used to divide the simulated gene expression values into 3’-UTR long and short isoform expressions, where we used Splatter ^18^ to simulate gene expression values.

On the simulated 3’-UTR isoform expressions, we compared scMAPA and scAPA. We did not consider the other methods because they cannot work in multiple cell groups. Across all *SD_isoprop_* values simulated for APA genes (0.06 to 0.18 around the value of the mouse data (0.127)), scMAPA consistently outperforms scAPA with higher sensitivity (**Fig. 2A**) while having similar specificity (**Fig. 2B**). While the above simulation fixed the number of APA and non-APA genes and the cell group sizes, we then ran other simulations by varying the number of APA and non-APA genes and the cell group size while fixing *SD_isoprop_* values for APA and non-APA genes (to 0.127 and 0.09, respectively). With various number of true APA genes (250, 500, and 1,000), scMAPA consistently outperforms that of scAPA in terms of sensitivity (**Fig. 2C and S. Fig. 2 C, E**) in all three group size distributions with a slight loss of specificity (**Fig. 2D and S. Fig. 2 D, F**).

**Figure 2.**
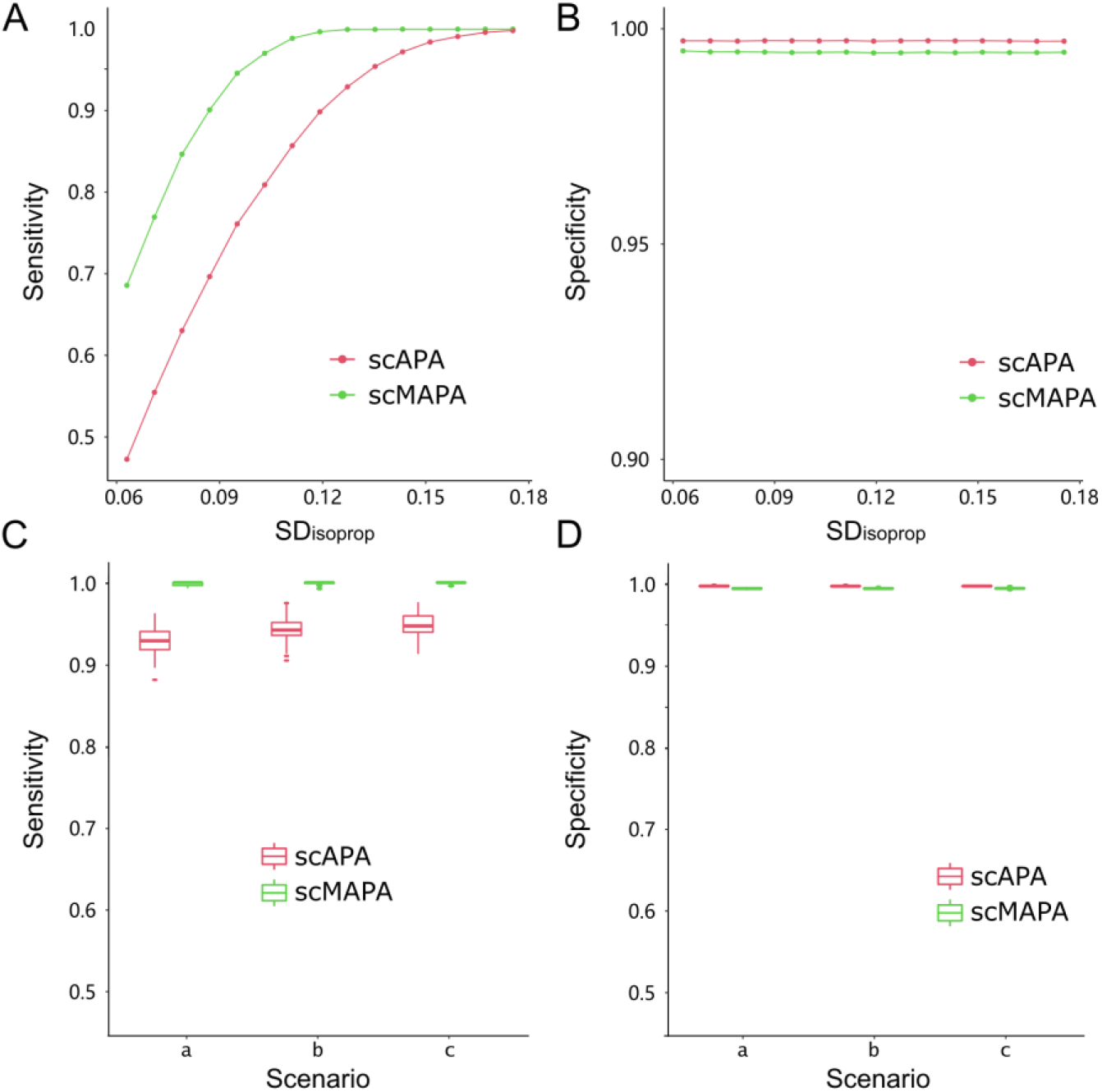
Performance assessment on the statistical component of scMAPA (regression + LRT) and scAPA (Pearson’s χ^2^) using simulated data. With fixed number of true APA events (500 out of 5000) and uniform distribution of cell cluster size (600 cells in each cell type), (A) sensitivity and (B) specificity were plotted against varying degree of standard deviation (SD) of PDUI values across clusters (SD_isoprop_) for true APA genes. With fixed number of true APA events (500) and SD values (0.127 for true APA genes and 0.009 for non-APA genes), (C) sensitivity and (D) specificity in scenarios with different distributions of cell cluster size: (20%, 20%, 20%, 20%, 20%) for scenario a, (30%, 17.5%, 17.5%, 17.5%, 17.5%) for b, and (50%, 12.5%, 12.5%, 12.5%, 12.5%) for c.

### scMAPA identification is accurate and robust

Further, we compared scMAPA with scAPA and Sierra using real biological data sets to show accuracy and robustness of scMAPA. scDAPA was not included in this comparison, because it does not return results that are compatible for the comparison, such as pA peaks, sites, or intervals. In the three PBMC data sets with various numbers of cells (1k, 5k and 10k data representing the number of cells), we defined various numbers of cell types (6, 8 and 13 types respectively) based on Seurat’s graph-based clustering ^19^ and annotated their cell types based on established marker genes ^20^ (see Methods, **S. Table 1**). We checked the proportion of the pA sites in proximity to the known pA sites annotated in PolyASite 2.0^21^ across various degrees of proximity. The higher the proportion is, the more of the method’s identified APA genes occurred close to the annotated pA sites. In the 10k and 5k data, scMAPA identifies the highest portion (**Fig. 3A**, **S. Fig. 3A, B**), showing the accuracy of scMAPA in determining pA sites.

**Figure 3.**
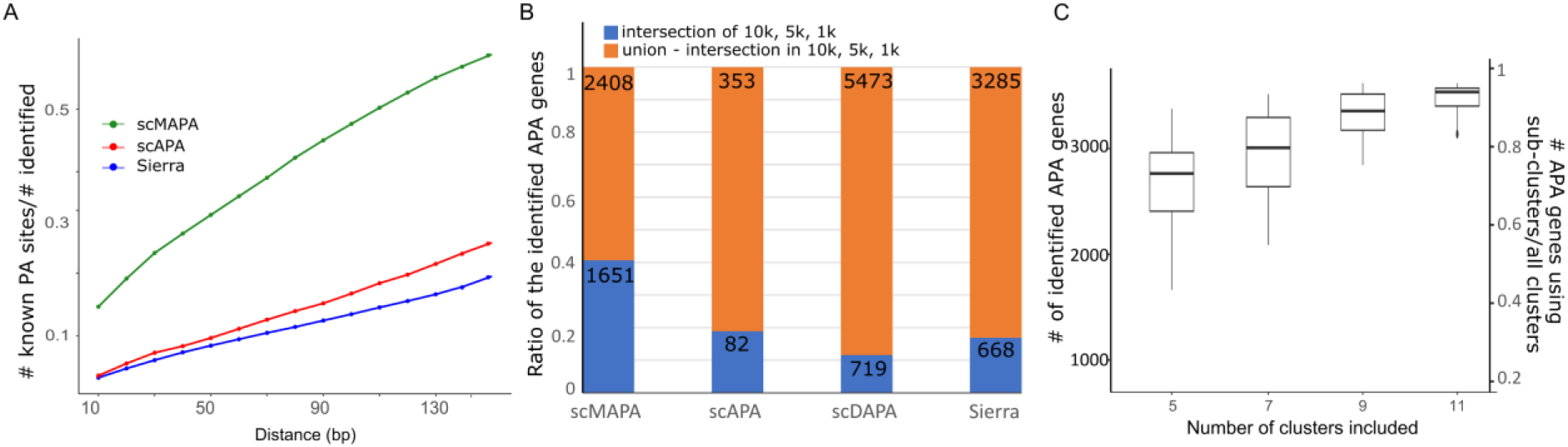
Performance assessment of scMAPA and scAPA using PBMC data. (A) Percentage of pA sites each method identified in the 10k data that are in proximity to known pA sites annotated in PolyASite 2.0 by the distance defining the proximity. (B) Ratio of significant APA genes found in all three PBMC data (10k, 5k, and 1k) in blue bar and in any combination but all three in orange by each method (C). Boxplot representing the number of APA genes identified by scMAPA by the number of clusters sampled from the 13 clusters of the 10k data.

We further evaluated the robustness of scMAPA in two ways. First, for each method, we identified APA genes for the PBMC data of various cell numbers (10k, 5k, and 1k). Across the data sets, scMAPA consistently identified similar sets of APA genes. On the other hand, scAPA, scDAPA and Sierra identified about half the size of APA genes across the data sets (**Fig. 3B**). Since the 1k, 5k, and 10k data are composed of a similar set of cell types from healthy adults (**S. Table 1**), the APA genes are expected to overlap across the three data sets. Thus, scMAPA identifying similar sets of APA genes across the data sets suggests that scMAPA identification is robust to the number of cells. Second, to show that scMAPA can robustly identify APA genes regardless of the number of cell types, we selected various numbers of cell types (5, 7, 9, and 11) from the 13 cell types in the 10k data. And, for each of the selections, we evaluated how many APA genes that were identified in the 13 cell types are recovered. Based on 50 random selections of various numbers of cell types (5, 7, 9, and 11), scMAPA is robust to the number of cell types (**Fig. 3C**). For example, when 5 cell types were sampled, 70.4% of all the APA genes were identified.

To demonstrate biological implications, we ran Ingenuity Pathway Analysis (IPA) on 1,432 APA genes identified only by scMAPA that are not identified by any other methods (**S. Table 2**). We checked that the genes indeed show various ratio of 3’UTR long and short isoforms across the clusters. For example, as FLT3 clearly showed a various usage of pA sites across the clusters (**S. Fig. 1 A, B**), it is included in the 1,432 scMAPA-unique APA genes. Further, GATA2 also showed various pA usages across the clusters and is included in the scMAPA-unique APA genes (**S. Fig. 1 C, D**). Interestingly, GATA2 was polyadenylated in the scRNA-Seq data of bone marrow mononuclear cell from acute myeloid leukemia patients^22^. Due to the developmental relationship between bone marrow and peripheral blood, GATA2 can undergo APA events also in the PBMC using similar molecular mechanisms. Collectively, the scMAPA-unique APA genes are significantly enriched (B-H p-value < 0.05) for multiple IPA Cellular Growth terms with implication for hematology developmental processes, including 9 with keyword “hemato” or “blood”. As “hemato” terms refer to diverse developmental processes of hematopoietic progenitor cells, previous reports on the role of APA in the hematopoietic stem cell differentiation^23^ supports the use of scMAPA. Altogether, scMAPA enables accurate and robust identification of dynamic APA in complex tissues.

### scMAPA identifies APA genes specific to each cell type

A novel function of scMAPA is to identify APA genes specific to each cell type (cell-type-specific APA genes) in the multi-group setting. To demonstrate this function in complex tissues, we analyzed the mouse brain scRNA-Seq data consisting of five major cell types: neurons, astrocytes, immune cells, oligodendrocytes and vascular ^17^ (see Methods). scMAPA identified 438 APA genes in neurons, 891 in immune, 374 in astrocyte, 422 in vascular and 430 in oligos, with some overlaps (**Fig. 4A**). Significant numbers of APA genes (35.4% on average, p-value<2.2e^−16^ by hypergeometric test) are differentially expressed (see Methods, **Fig. 4B**). Since APA genes are more likely differentially expressed than non-APA genes (supplemental material)^24,25^, this result suggests that scMAPA identified APA genes are indeed functional APA genes.

**Figure 4.**
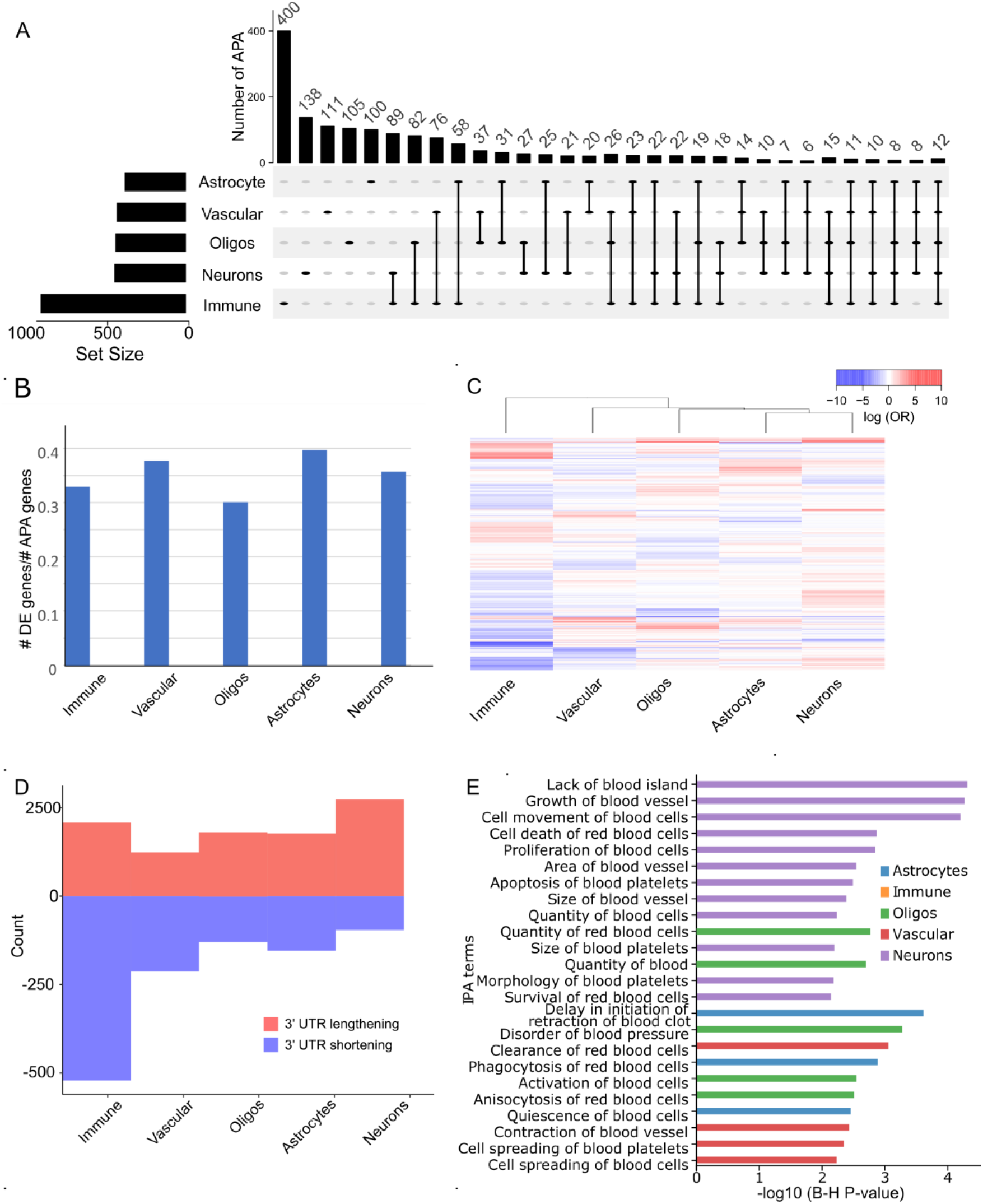
A novel module of scMAPA Cluster-specific APA identification on mouse brain data. (A) Upset plot showing APA genes specific to each cell type. (B) Ratio of DE genes in the APA genes specific to each cell type. (C) Heatmap of coefficients of cell type-specific APA genes. Coefficients were estimated in logistic regression model. (D) Bar plot shows the number of 3’-UTR lengthening and shortening detected in each cell type. (E) Bar plot shows the enrichment (-log10(B-H p-value)) of brain cell-type-specific APA genes (blue for astrocyte, orange for immune, green for oligos, red for vascular, and violet for neurons). Only the highest bar for each term is displayed.

For each cell type, we further investigated which genes undergo APA events in which direction (shortening/lengthening) and to which degree. For each APA gene, the degree was estimated by the regression coefficients and the direction of APA events was estimated by the sign of the coefficients (see Methods). First, by implementing hierarchical clustering to the coefficients (**Fig. 4C**), scMAPA found that immune and neuron cells are most distinguished from the other cell types, systematically confirming the previous finding of scAPA that immune and neuron cells are most different in the APA pattern ^26^. Second, by identifying the direction of the significant APA events (shortening/lengthening) by cell type, scMAPA found that neuron cells are characterized with 3’-UTR lengthening (**Fig. 4D**), which is consistent with previous findings of the dominance of 3’-UTR lengthening in neuron cells ^27–30^. Third, by running IPA on the 438 neuron-specific APA genes, scMAPA brings insights to identify the functional role of APA on the neuron-specific biological functions. For example, the neuron-specific APA genes (**S. Table 5)** are significantly (B-H p-value < 0.05) and exclusively enriched for the neuron-specific biological terms, including 11 blood or blood vessel disease terms, such as Proliferation/Survival of blood cells and Area/Size of blood vessel (**Fig. 4E, S. Fig. 4**). Neuron and blood cells interact to allow ready exchange of nutrients and waste products, enabling the high metabolic activity of the brain despite its limited intrinsic energy storage^31^. Although this interaction is believed to play critical function in maintaining the operational condition of brains, little is known as to how this highly dynamic process is tightly regulated. Our findings on the neuron-specific APA genes suggest that APA contributes to the tight regulation of this intricate biological mechanism. Together, scMAPA’s cell-type-specific APA genes facilitate investigating functional implications of APA in a cell-type-specific manner.

### sciMAPA controls confounding factors

To demonstrate how scMAPA controls confounders, we will identify cell-type-specific APA genes and brain regions as confounder. For this, we first split the mouse brain scRNA-Seq data by both cell type (neurons, immune cells, astrocytes, oligos, and vascular cells) and brain region (cortex and midbrain dorsal) information that is annotated^17^. Since the cell type and the brain region are independent to each other (**Fig. 5A, B**), we quantified 3’-UTR long and short isoforms in each combination (5 cell types ×2 brain regions) using scMAPA. To first show how brain region information confounds cell-type-specific APA analysis, we set scMAPA in two different runs: one where only the cell type is set as covariate and the other where the cell type is as covariate and brain region is as confounder. In the model with the cell type covariate and the brain region confounder, 113 APA genes were excluded as compared to the model only with the cell type covariate (**S. Table 6**). To confirm that the 113 genes play roles specific to the brain regions, we investigated whether the genes are up-regulated in the Genotype-Tissue Expression (GTEx)^32^ brain and non-brain tissues (see Methods). Indeed, the genes are up-regulated in brain samples than in non-brain (p-value=6.57e^−105^, **Fig. 5C**). In addition, we checked whether the 113 genes are especially up-regulated in brain cortex samples than brain non-cortex samples (p-value=5.86e^−7^, **Fig. 5D**). Since the 113 genes are not enriched in down-regulated genes in GTEx tissues (**S. Fig. 5A, B**). On the other hand, when we investigated 2,715 APA genes that are identified when both the cell types and brain region are used as covariates, we found that these genes are up-regulated in brain tissues, but not necessarily in cortex tissues (**S. Fig. 5C**, **D**). Ingenuity Pathway Analysis (IPA) upstream regulator analysis on the 113 genes (**S. Table 7**) further validates the relevance of 113 APA genes to brain cortex. For example, IL1B, LRRC4, and TREX1 are three of the most significant upstream regulators of the 113 genes. All three genes are known to express specifically in the cortex region, heavily involved in the development of brain cortex ^33–35^. The results suggest that scMAPA can control brain regions as confounders in identifying APA and remove confounders that may conflict functional analyses on the cell-type-specific roles of APA.

**Figure 5.**
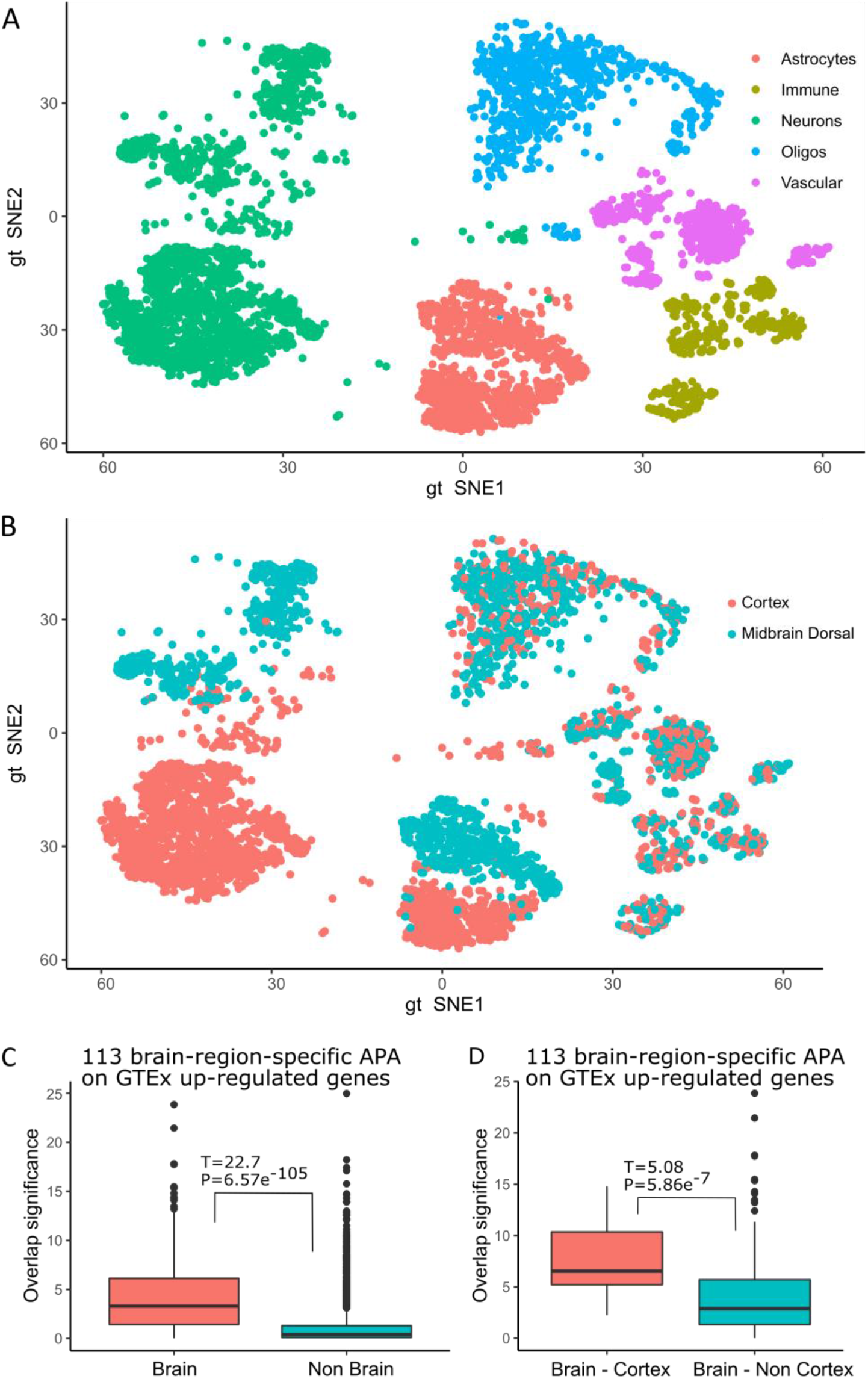
tSNE plot showing the cell type (A) and brain region (B) of the mouse brain scRNA-Seq data. (C) Significance of overlap between the 113 brain-region-specific APA genes and the up-regulated genes in GTEx samples whether they are from brain (red) or not (green). A higher overlap significance indicates a more significant overlap, calculated by Enrichr. (D) Significance of overlap between the 113 genes and the up-regulated genes in GTEx brain samples whether they are from cortex (red) or not (green).

## DISCUSSION

To identify APA genes in scRNA-seq data, we developed scMAPA that is novel in several ways. First, while existing methods operate with assumptions on the shape of input scRNA-Seq data, scMAPA quantifies 3’-UTR long and short isoforms without posing such assumptions. Thus, scMAPA enables an accurate and robust APA identification by considering genes that may not follow the assumptions of the existing methods. As a result, scMAPA outperforms existing methods in identifying APA genes in various simulation (**Fig. 2**) and the PBMC data (**Fig. 3**) while it returns consistent results with existing methods (see Supplemental Material). As the second novelty, it identifies APA genes specific to each cell type using a sophisticated statistical model, enhancing interpretability on the APA genes. This novel analytical layer further elucidates cell-type-specific function of APA in the mouse brain data (**Fig. 4, 5**). For example, APA genes unique to neuron cells suggest the potential role of APA for the interaction between neurons and blood vessels, which is critical to maintain the operational condition of brains. The sophistication of the model helps control confounding factors by allowing to incorporate confounders directly in the model. In our cell-type-specific APA identification in the mouse brain data with brain region as the confounder, scMAPA can factor out the 113 APA genes that are likely related to a specific brain region (brain cortex). By factoring out such biological signal that is not likely specific to any brain cell type, scMAPA can accurately conduct functional analyses on the cell-type-specific roles of APA. Lastly, we developed a simulation platform based on the distribution of 3’-UTR long and short isoforms in which to assess performance of APA identification methods. This simulation platform enables to not only compare performance of APA identification methods, scMAPA and scAPA, but also break down their performance by 3’-UTR isoform distribution across cells.

Especially, it is important to note that scMAPA makes point estimations of the pA sites. Although point estimations are more directly relevant than interval estimations for further analyses, e.g. conducting omics analyses and designing experiments, point estimation methods are generally disadvantageous in checking the distance with the annotated pA sites (**S. Fig. 3A** and **S. Fig. 3B**), because it returns a single point to calculate the proximity while interval estimation returns two points (start and end of the interval). Still, scMAPA outperforms the interval estimation results of Sierra and scAPA, while the interval estimation results are better than point estimation results of Sierra and scAPA (**S. Fig. 3A, B**). Also, it is worth noting that scMAPA identifications are consistent with the results of the other methods. After identifying APA genes in multi-group settings, with minimal modifications for scDAPA and Sierra (see Methods), scMAPA identifies an intermediate number of APA genes between scDAPA and Sierra/scAPA (10k in **Fig. 3C** and 5k in **S. Fig. 3D**), more than half of the scMAPA’s findings are found in other methods (59.9% for 10k and 51.9% for 5k). This shows that, although scMAPA is the only method designed with an optimization criterion, rather than with signal processing steps, this high overlaps with other methods validate the use of scMAPA.

With the improved accuracy/robustness and enhanced interpretability, scMAPA is extendible in the following directions. First, while scMAPA assumes two types of 3’-UTR isoforms: long and short, and assumes no pA sites on the introns, recent works reported genes with more than two 3’-UTR isoforms^21^ and pA sites on the introns^3^. With these studies, scMAPA can be extended to incorporate such cases. Second, as we transform 3’-biased reads to represent full-length 3’UTR of the transcripts, this transformation allows to apply other established methods developed for bulk RNA-Seq data, which represent full-length transcripts, such as APATrap^36^, TAPAS^37^, and DaPars^24^ for scRNA-Seq data analysis. Third, due to this transformation, scMAPA is directly amenable for other scRNA-Seq data that are not 3’tag-based (e.g. Smart-seq2^38^). scMAPA is also applicable for bulk RNA-Seq data sets that are collected from multiple biological conditions.

Altogether, we developed a statistical method to identify APA genes in the multi-group setting. With high sensitivity and interpretability, scMAPA allows to understand cell-type-specific function of APA events, which is essential to shed novel insights into the functional roles of dynamic APA in complex tissues.

## Acknowledgements

We thank Daniel Weeks, Ph.D., Professor, Department of Human Genetics, University of Pittsburgh for valuable discussion. This research was supported in part by the University of Pittsburgh Center for Research Computing through the resources provided. We also acknowledge the authors of scAPA for their generous provision of their data.

## Author Contributions

H.J.P and Y.B. conceived the project, designed the experiments. Y.B. and Z.F. implemented the software. Y.B., Y.Q., R.M. performed the analysis. S.K., K.N., H.M.Z., R.K., Q.P. interpreted the results statistically and/or biologically.

## Competing interests

The authors declares no competing financial interests.

## Availability of data and materials

The open source scMAPA program (version 0.9.1) is freely available at https://github.com/ybai3/scMAPA with necessary example data for this analysis.

## Funding

This work was supported partly by the Joan Gollin Gaines Cancer Research Fund at the University of Pittsburgh to H.J.P.. This project used the UPMC Hillman Cancer Center Biostatistics Shared Resource that is supported in part by award P30CA047904.

**S. Table 1.** Cell type annotation based on marker genes curated in CellMarker^20^ for 10k, 5k, and 1k in the PBMC data.

**S. Table 2.** Detailed information of APA genes detected by scMAPA, scAPA, scDAPA, and Sierra on the PBMC data including Ingenuity Pathway Analysis (IPA) analysis result.

**S. Table 3.** scMAPA estimation result for cell-type-specific APA genes on the mouse brain data.

**S. Table 4.** Result of IPA comparison analysis on the “Disease & Function” terms enriched for APA genes identified uniquely by scAPA, scMAPA and commonly by both on the mouse brain data (1,446, 2,175, and 1,048 respectively).

**S. Table 5.** Result of IPA comparison analysis on the “Disease & Function” terms enriched for APA genes identified uniquely in astrocyte, immune, oligos, vascular, and neuron cells.

**S. Table 6.** scMAPA estimates on the input data that are split by cell type and brain region either with brain region as a confounder or not.

**S. Table 7.** IPA upstream regulator analysis result (enrichment p-value) on 113 and 2,715 APA genes that are supposed to be brain-region-specific and non-specific, respectively.

## Notes

http://mousebrain.org/

https://support.10xgenomics.com/single-cell-gene-expression/datasets/3.0.0/pbmc_10k_v3

### Competing Interest Statement

The authors have declared no competing interest.

