## Supplemental Figures for "Cell-type-specific alternative polyadenylation (APA) genes reveal the function of dynamic APA in complex tissues"

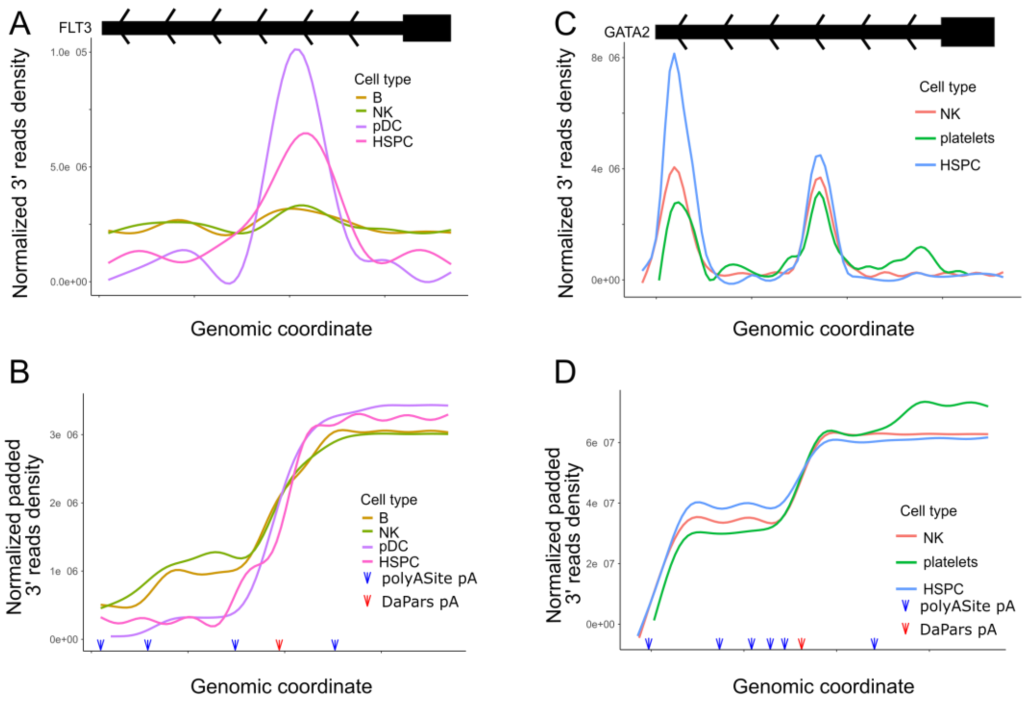


E

| | scMAPA | scAPA | scDAPA | Sierra |
|---|---|---|---|---|
| **Step 1.** split reads by clusters | In Python script | Dropseq-tools, called in R | In shell script | User provided |
| **Step 2.** quantify APA in each cluster | DaPars + long/short isoform estimation | Homer findPeaks (peak filtering) + mclust (splitting peaks) + featureCounts (isoform estimation) | Based on difference in read length and in peak distribution | Junction awared peak calling<br>Peaks merging for multiple datasets<br>UMI counting |
| **Step 3.** estimate significance of APA across clusters | Binary or multinomial logistic regression + LRT | Pearson's $\chi^2$ test | Wilcoxon sum-rank test on pairwise comparisons | DEXSeq pipline on pseudo bulk counts |
| **Step 4.** define cluster-specific APA | Wald test + filter on estimated coefficients | | | |
| Note | Identifying up to 2 peaks | Identifying > 2 peaks | Identifying up to 2 cell clusters | Identifying up to 2 cell clusters |

**Supplemental Figure 1.** Signal density of 10k PBMC scRNA-Seq reads mapped onto 3′-UTR of GATA2 (A) and FLT3 (B) in terms of original 3′ tag-based (top panel) or of padded reads (C) and (D) respectively, for selected clusters for presentation purpose. While the genomic coordinates are shared between A and C, B and D, the blue arrows indicate the polyA site annotated in polyASite database (v. 2.0). Red arrow indicates the proximal polyA site predicted. (E) Algorithm overview of bioinformatic tools and statistical methods to identify dynamic APAs in scRNA-Seq data

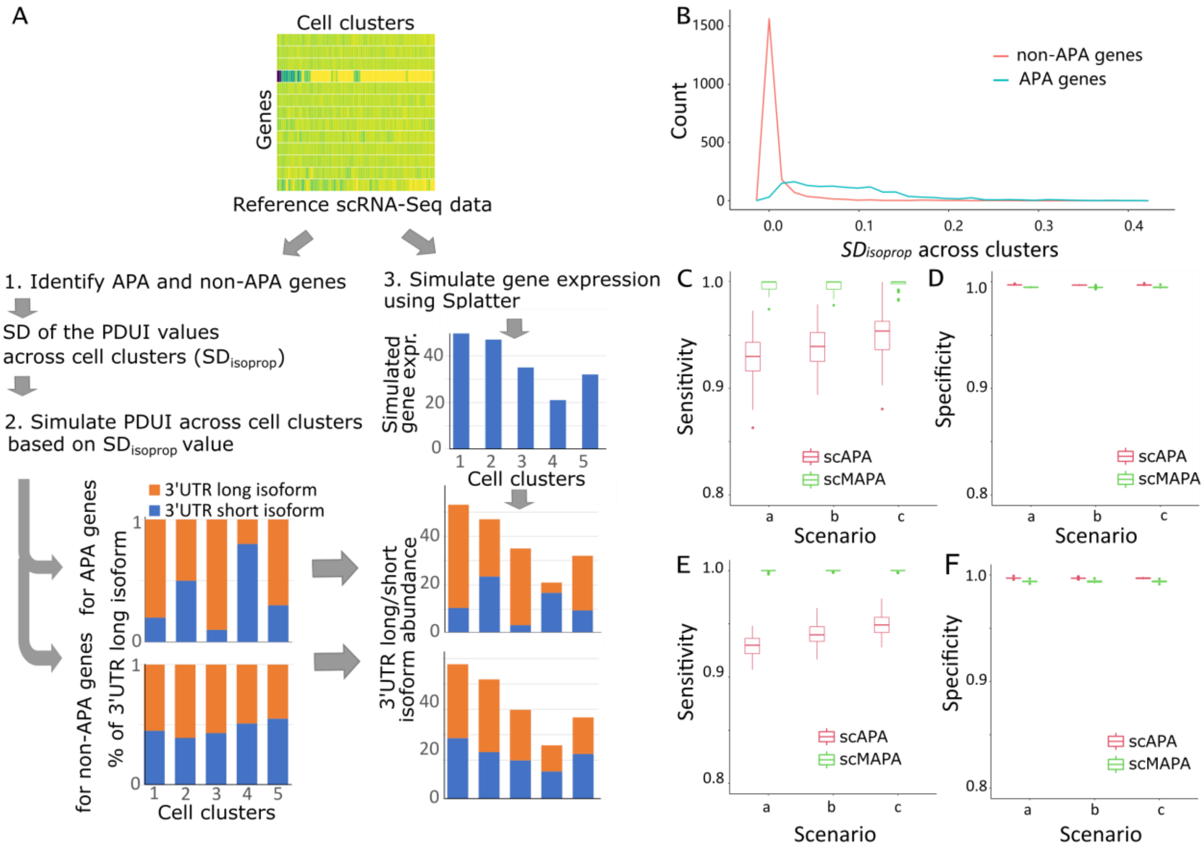


**Supplemental Figure 2**. Performance assessment on the statistical component of scMAPA (regression + LRT) and scAPA (Pearson's $\chi^2$) using simulated data. (A). Illustration of the simulation process. Genes identified as significant APA genes by both scMAPA and scAPA were considered as APA genes. Genes identified as non-significant APA genes by both methods were considered as non-APA genes. (B) shows the frequency of standard deviations (SD) of PDUI values across clusters from mouse brain data. (C) to (F) show the performance assessment using simulated data. With fixed number of true APA events (250) and SD values (0.1268 for true APA genes and 0.009190 for non-APA genes), box plots in (C) and (D) show the sensitivity and specificity in scenarios with different distributions of cell type populations: (20%, 20%, 20%, 20%, 20%) for scenario a, (30%, 17.5%, 17.5%, 17.5%, 17.5%) for b, and (50%, 12.5%, 12.5%, 12.5%, 12.5%) for c. Box plots in (E) and (F) show the sensitivity and specificity with the number of true APA events set to 1,000 and all other factors remain same.

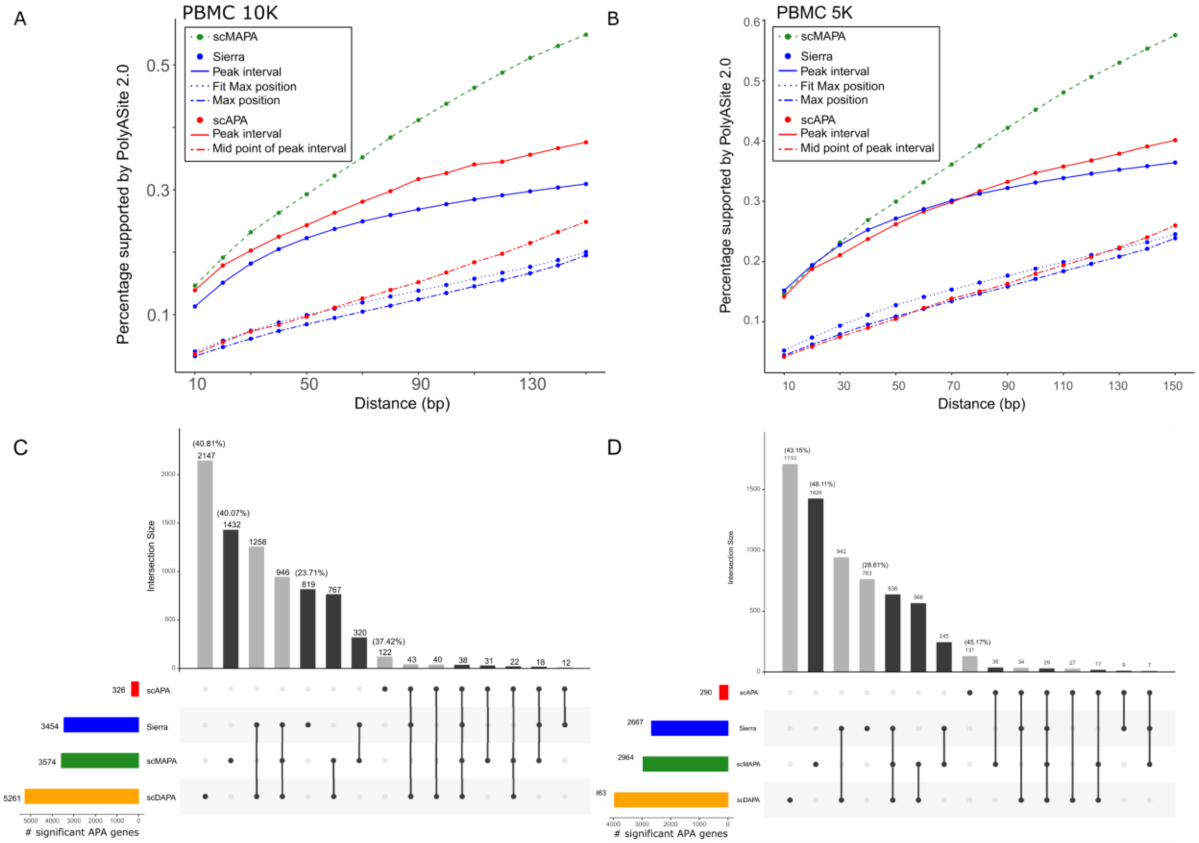


**Supplemental Figure 3.** Percentage of pA sites each method identified in the PBMC 10k (A) and 5k (B) data that are in proximity to known pA sites annotated in PolyASite 2.0 by the distance defining the proximity. Upset plot showing diverse overlaps among APA genes in the 10K (C) and 5k (D) data identified by four methods, scAPA, Sierra, scMAPA and scDAPA. Barplot on top shows the number of genes corresponding to the set combination indicated below. Black bars correspond to the sets involving scMAPA results. Colored horizontal bars represent the total number of APA genes identified by each method.

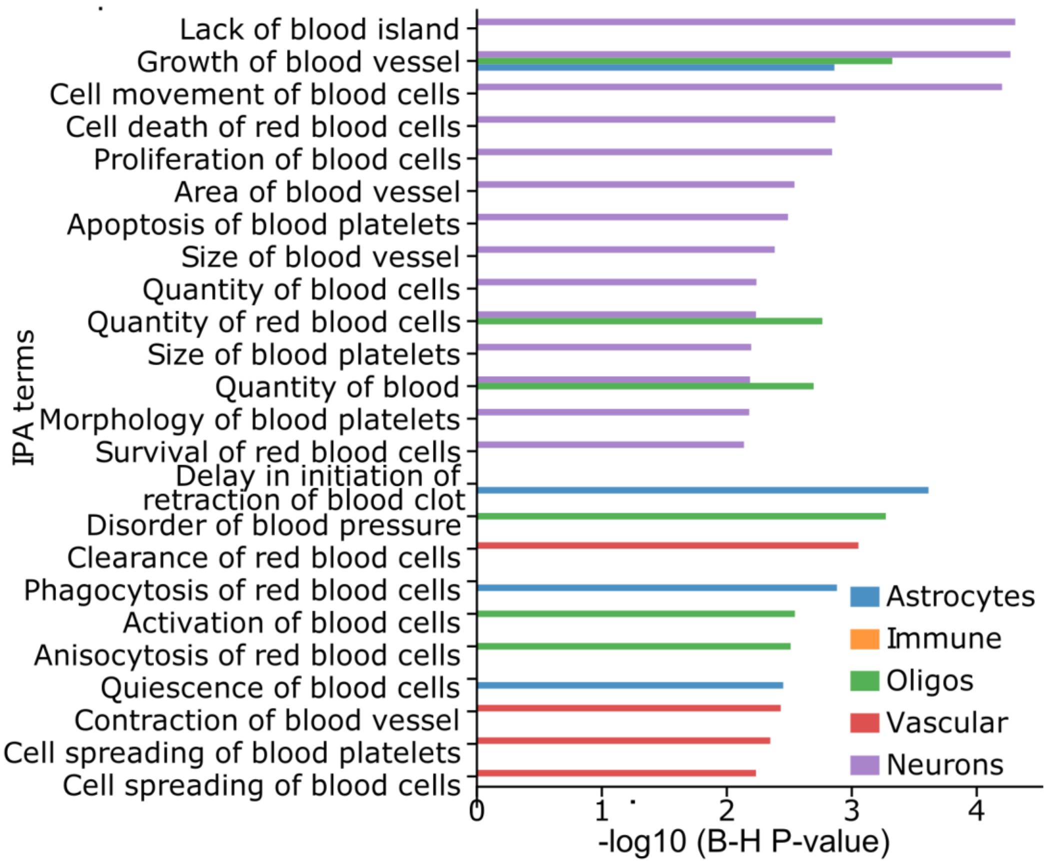


**Supplemental Figure 4.** Bar plot shows the enrichment (-log10(B-H p-value)) of brain cell-type-specific APA genes (blue for astrocyte, orange for immune, green for oligos, red for vascular, and violet for neurons). All significant terms (B-H p-value < 0.01) are displayed.

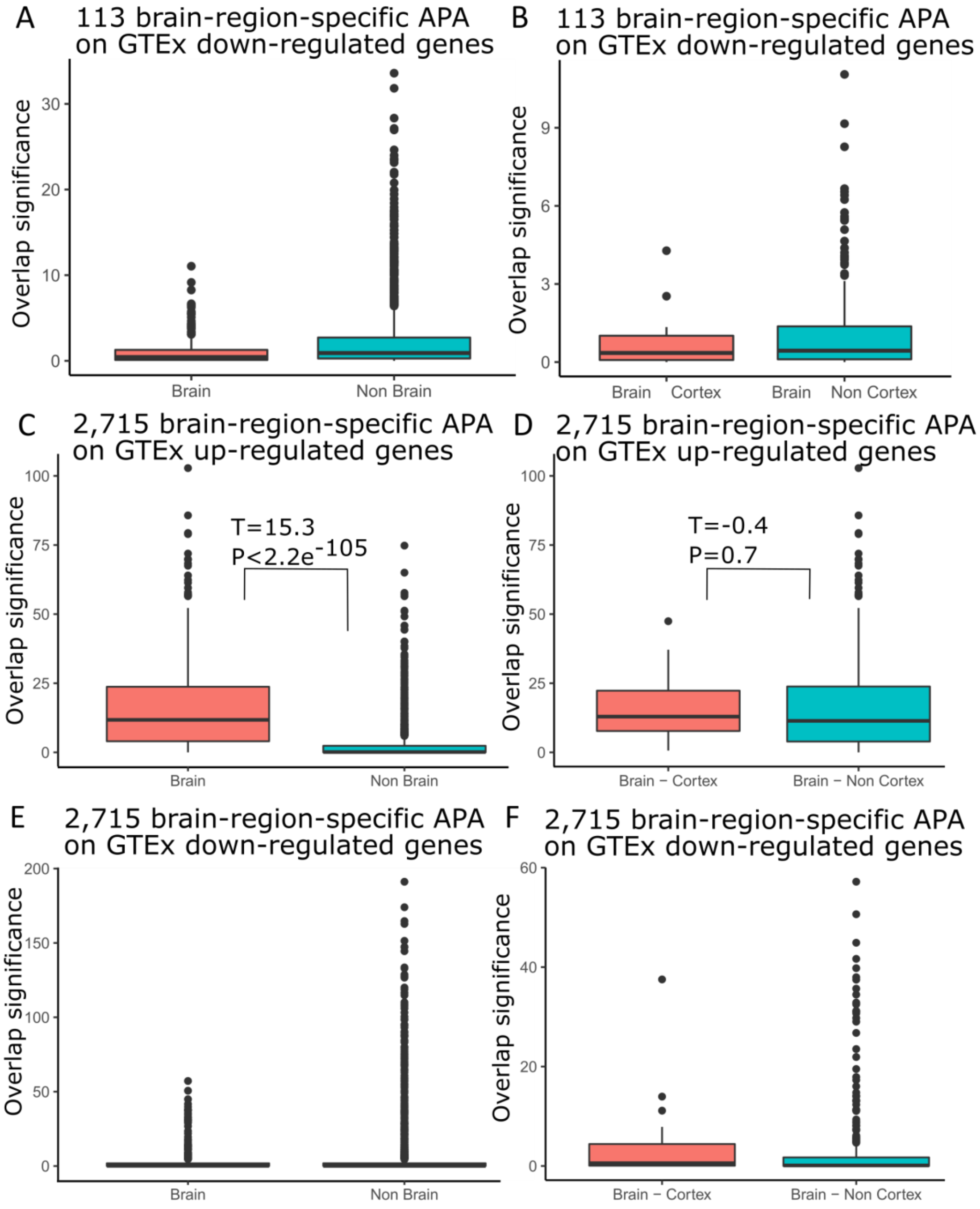


**Supplemental Figure 5.** Significance of overlap between the 113 brain-region-specific APA genes and the down-regulated genes in GTEx samples (A) whether they are from brain (red) or not (green) (B) whether they are from brain cortex region (red) or not (green). Significance of overlap between the 2,715 APA genes that are not specific to brain regions and the up-regulated genes in GTEx samples (C) whether they are from brain (red) or not (green) (D) whether they are from brain cortex region (red) or not (green). Significance of overlap between the 2,715 APA genes that are not specific to brain regions and the down-regulated genes in GTEx samples (C) whether they are from brain (red) or not (green) (D) whether they are from brain cortex region (red) or not (green). A higher overlap significance indicates a more significant overlap, calculated by Enrichr.
